# Popcorn haploids identified by Navajo phenotype and ploidy level

**DOI:** 10.1101/2023.03.16.533026

**Authors:** Jean Paulo Aparecido da Silva, José Marcelo Soriano Viana, Kaio Olimpio das Graças Dias, Jéssica Coutinho Silva, Vivian Torres Bandeira Tupper, Wellington Ronildo Clarindo

**Affiliations:** Department of General Biology, Federal University of Viçosa, Viçosa, Brazil

**Keywords:** *Zea mays* L. var. Everta, haploid induction, doubled haploid, unilateral cross incompatibility, Genetic improvement

## Abstract

For popcorn, obtaining and identifying haploids are still challenging steps. We aimed to induce and screen haploids in popcorn using the Navajo phenotype, seedling vigor and ploidy level. We used the Krasnodar Haploid Inducer (KHI) in crosses with 20 popcorn source germplasms and five maize controls. The field trial design was completely randomized, with three replications. We assessed the efficacy of induction and of identification of haploids based on haploidy induction rate (HIR) and false positive and negative rates (FPR and FNR). Additionally, we also measured the penetrance of the Navajo marker gene (*R1-nj*). All putative haploids classified by *R1-nj* were germinated together with a diploid sample and evaluated for false positives and negatives based on vigor. Seedlings from 14 females were submitted to flow cytometry to determine the ploidy level. The HIR and penetrance were analyzed by fitting a generalized linear model with a logit link function. The HIR of the KHI, adjusted by cytometry, ranged from 0.0 to 1.2%, with a mean of 0.34%. The average FPR from screening based on the Navajo phenotype were 26.2% and 76.4%, by the vigor and ploidy, respectively. The FNR was zero. The penetrance of *R1-nj* ranged from 30.8 to 98.6%. The average number of seeds per ear in temperate germplasm (76) was lower than that obtained in tropical germplasm (98). There is induction of haploids in germplasm of tropical and temperate origin. We recommend the selection of haploids associating the Navajo phenotype with a direct method of confirming the ploidy level, such as flow cytometry. We also show that haploid screening based on Navajo phenotype and seedling vigor reduces misclassification. The origin and genetic background of the source germplasm influence the *R1-nj* penetrance. Because the known inducers are maize, developing doubled-haploid technology for popcorn hybrid breeding requires overcoming the unilateral cross-incompatibility.

## Introduction

The doubled-haploid (DH) technology allows for obtaining homozygous genotypes in a shorter period of time, increasing genetic gains and efficiency in hybrid development, associated with lower costs (Melchinger et al., 2005; Chaikam et al., 2019; Molenaar et al., 2019). The development of DH lines consists of the induction of seeds with haploid embryos, identification of these seeds, duplication of the chromosome set, and multiplication of DH_0_ seeds (Prigge and Melchinger, 2012). Haploid induction is based on the *in vivo* induction method, which consists of using haploid inducers in crosses with a germplasm source. These inducers can be used as pollen donors, generating gymnogenetic haploids, or as pollen receptors, generating androgenetic haploids (Sarkar and Coe, 1966). Inducers are usually homozygotes for the *R-navajo gene* (*R1-nj*), a dominantly acting marker that causes purple coloration of the scutellum and aleurone layer (Navajo phenotype). Scutellum and aleurone layer staining is used as embryo and endosperm markers, respectively, to identify haploid seeds (Kleiber et al., 2012).

The haploid induction rate (HIR) is the most important characteristic of a haploid inducer. The HIR is calculated by the ratio of the number of haploid seeds to the total number of seeds produced in an induction cross and is the common parameter to compare the haploid induction efficiency of different inducers (Almeida et al., 2020). This characteristic is influenced by the genetic background inducer and by the source germplasm (Chase, 1952; Coe, 1959; Lashermes and Beckert, 1988; Eder and Chalyk, 2002; Prigge et al., 2011). Currently, induction rates vary on average from 8-15% (Röber, 1999; Rotarenco et al., 2010; Chaikam et al., 2019). *In vivo* haploid induction is a quantitative trait controlled by a large number of unknown loci (Lashermes and Beckert, 1988; Röber et al., 2005). It was found that the *qhir11* subregion has a significant effect on the induction rate (Nair et al., 2017). Within this region, a gene encoding *phospholipase A* was identified as responsible for haploid induction called *MATRILINEAL* (*MTL*) (Kelliher et al., 2017; Liu et al., 2017; Almeida et al., 2020).

The *C2* or *Whp1, A1*, and *A2* genes are required for anthocyanin biosynthesis. Homozygous recessive genotypes for any of these genes do not synthesize anthocyanin and the absence of a dominant allele interrupts the pathway in the aleurone layer (Ford, 2000). The variation in *R1-nj* expression, both for the labeled area and for the intensity of anthocyanin in the endosperm and embryo, affects the accuracy of haploid identification. The low expression of the marked area and the low intensity of anthocyanin staining in the endosperm or embryo are generally associated with high rates of misclassifications (Prigge et al., 2011). In a haploidy-inducing cross, depending on the homozygosity and heterozygosity of the inhibitory alleles in the population, the *R1-nj* marker may be completely inhibited or color expression may segregate (Chaikam et al., 2015). The advantage of *R1-nj* is the possibility of identifying haploid individuals even before germination.

Fertilization in maize occurs through the interaction between the male gametophyte (pollen) and the female sporophyte (silk). The *in vivo* induction of haploids strictly depends on this process. However, the existence of an incompatibility system that affects fertilization between some maize and popcorn genotypes can prevent the occurrence of fertilization and, consequently, the induction of haploidy. The main incompatibility system known in maize is the *gametophyte factor 1* (*ga1*) (Kermicle and Evans, 2010; Zhang et al., 2018). For this system, three alleles are known, *ga1, Ga1-s*, and *Ga1-m*. The *ga1* allele is recessive and receptive to any pollen genotype, but cannot pollinate *Ga1-s* plants (Kermicle and Evans, 2005). The *Ga1-s* allele is codominant, receptive to *Ga1-s* and *Ga1-m* pollen, but not to *ga1* pollen (Demerec, 1929; Schwartz, 1950; Kermicle, 2006). The *Ga1-m* allele is dominant and capable of pollinating and being pollinated by any of these alleles (Mangelsdorf and Jones, 1926; Nelson, 1952). The *Ga1-s* allele is common in popcorn (Kermicle, 2006), while most other types of corn, including the known inducers, carry the *ga1* allele and therefore are susceptible to the fertilization barrier imposed by *Ga1-s* (Moran Lauter et al., 2017).

Although the DH technology could potentially accelerate maize reproduction, it has been little explored in the development of popcorn lines. In popcorn, there is no information about the basic steps such as induction and haploid identification. Therefore, our objective was to induce *in vivo* maternal haploids and identify them by Navajo phenotype, seedling vigor, and ploidy level.

## Materials and methods

### Genetic material

The Krasnodar haploid inducer (KHI), a temperate population, was used as the gymnogenetic inducer (Milani et al., 2016). The females included eight popcorn populations, 10 popcorn inbred lines, seven popcorn interpopulation hybrids, three maize single crosses, a maize two-way cross, and a maize population (Table 1). The tropical populations Viçosa C4 and Beija-Flor C4 were obtained from the Viçosa and Beija-Flor populations, respectively, after four half-sib selection cycles. The tropical population Synthetic UFV was derived by crossing 40 elite inbred lines from Viçosa and Beija-Flor. The five temperate populations (UFV-MP1, UFV-MP2, UFV-MP3, UFV-MP4, and UFV-MP5) were derived from the North American hybrids P622, P625, P802, AP2501, and AP4502, respectively, developed by the Agricultural Alumni Seed Improvement Association, Romney, IN, USA. The inbreed progeny 18-303-2 and 18-390-1 were derived from UFV-MP1, 19-206-6 from UFV-MP2, 18-176-5 from UFV-MP3, 19-504-4 from UFV-MP5, 20-2039 and 20-2064 from Viçosa C4 and 20-2009, 20-2010 and 20-2024 were derived from Beija-Flor C4.

**Table 1.** Sources of germplasm used in haploidy induction crosses with the KHI.

| Germplasm | Genetic basis | Origin | Type |
| --- | --- | --- | --- |
| 19-1206-6 | Inbred line | Temperate | Popcorn |
| 19-1504-4 | Inbred line | Temperate | Popcorn |
| 18-390-1 | Inbred line | Temperate | Popcorn |
| 18-303-2 | Inbred line | Temperate | Popcorn |
| 20-2039 | Inbred line | Tropical | Popcorn |
| 18-176-5 | Inbred line | Temperate | Popcorn |
| 20-2009 | Inbred line | Tropical | Popcorn |
| 20-2010 | Inbred line | Tropical | Popcorn |
| 20-2064 | Inbred line | Tropical | Popcorn |
| 20-2024 | Inbred line | Tropical | Popcorn |
| UFV-MP1 | Population | Temperate | Popcorn |
| UFV-MP2 | Population | Temperate | Popcorn |
| UFV-MP3 | Population | Temperate | Popcorn |
| UFV-MP4 | Population | Temperate | Popcorn |
| UFV-MP5 | Population | Temperate | Popcorn |
| Viçosa C4 | Population | Tropical | Popcorn |
| Beija-Flor C4 | Population | Tropical | Popcorn |
| Synthetic UFV | Population | Tropical | Popcorn |
| Viçosa C4 x Beija Flor C4 | Interpopulation hybrid | Tropical | Popcorn |
| Beija-Flor C4 x Synthetic UFV | Interpopulation hybrid | Tropical | Popcorn |
| UFV-MP2 x UFV-MP5 | Interpopulation hybrid | Temperate | Popcorn |
| Viçosa-C4 x UFV-MP2 | Interpopulation hybrid | Tropical x Temperate | Popcorn |
| Viçosa C4 x UFV-MP4 | Interpopulation hybrid | Tropical x Temperate | Popcorn |
| Viçosa C4 x UFV-MP5 | Interpopulation hybrid | Tropical x Temperate | Popcorn |
| Synthetic UFV x UFV-MP1 | Interpopulation hybrid | Tropical x Temperate | Popcorn |
| 2B710PW | Single-cross | Tropical | Maize |
| MG580PW | Single-cross | Tropical | Maize |
| ES481PW | Single-cross | Tropical | Maize |
| Feroz | Two-way cross | Tropical | Maize |
| AGS Lavrador OP | Population | Tropical | Maize |

### Induction crosses

The induction crosses occurred naturally in the 2021/2022 growing season in an isolated field located in an experimental area at Federal University of Viçosa (UFV) in the municipality of Viçosa (20°50’30”S, 42°48’30”W, at an altitude of 720 m). The experiment design was completely randomized, with 25 treatments and three replications. Each pollination plot consisted of two 5-m rows of females to be induced. They were detasseled before anther dehiscence. Four rows of the inducer were planted between each plot at four different times. The first planting was performed one week after sowing the females. The other plantings were performed at five-day intervals after the first inducer plantings. An outer border with two rows of the inducer was also sown. The females and the inducers were planted with a spacing of 20 cm between plants and 70 cm between rows. Thinning was performed at stage v3 to remove the less developed plants so that there were five plants per m.

### Identification of haploids

For each replication, the seeds resulting from the crosses were visually separated according to the appearance of purple on the endosperm and embryo (Navajo phenotype), based on the methodology described by Nanda and Chase (1966), to identify haploids; thus, seeds with purple endosperms and white embryos were classified as haploids, those with purple endosperms and purple embryos were classified as diploids, and the seeds without anthocyanin expression in the embryo and endosperm were classified as yellow. In addition to the classes proposed by Nanda and Chase (1966), the purple seed class was also considered a measure of *R1-nj* penetrance, which was defined by the sum of the haploid and diploid classes. Based on this classification, the estimates of the haploid induction rate (HIR) based on the *R1-nj* and gene penetrance were obtained by HIR = haploids/total and Penetrance = purples/total.

### Seedling vigor

To evaluate vigor, the putative haploid seeds obtained in each pollination plot were germinated in trays containing two sheets of germitest paper and filled with sand. The number of haploid seeds obtained in each plot was variable, so the number of seedlings evaluated in each replicate and treatment was different. In addition to the haploid seeds, 10 putative diploid seeds and 10 seeds of the germplasm from which they originated were also germinated. Before planting, all seeds were treated with the fungicide. The relative vigor of the seedlings was evaluated 10 days after sowing. The evaluation was performed using the putative diploids as a reference. The taller seedlings with greater diameter leaves and stems and purple coloration were classified as diploid.

### Ploidy level determination by flow cytometry

After evaluating vigor, the haploid seedlings from 14 of the 25 induced females, including those classified as false positives (FP) based on vigor, were subjected to flow cytometry to confirm the ploidy level. For each treatment, a sample from a diploid individual and a sample from the germplasm source were used as external standards. For nuclear extraction, fragments of young leaves of the evaluated individuals and the external standards were chopped in 500 µL of OTTO-I Otto (1990) buffer supplemented with 2.0 mM dithiothreitol (Sigma®). After 3 min, 500 µL of the same buffer was added, and the homogenate was filtered through 30 µm nylon mesh (Partec®) and centrifuged at 100 × g for 5 min. The supernatant was discarded, and the pellet was resuspended and incubated for 10 min in 100 μL of the OTTO-I buffer. The nuclear suspensions were stained in 500 µL of OTTO-I:OTTO-II buffer (1:2), supplemented with 2.0 mM dithiothreitol (Sigma®) and 75 µM propidium iodide (Sigma®) at room temperature for 30 min in the dark, filtered through 20 µm nylon mesh (Partec®) Praça-Fontes et al., (2011), and analyzed in a BD Accuri C6 flow cytometer. The histograms obtained from the nuclear suspensions of the possible haploid seedlings were compared to those provided by the external standards to determine the relative ploidy level of each sample.

### Statistical analysis

The putative haploids and putative diploids that were confirmed by the vigor test and flow cytometry were categorized as true positives (TP) and true negatives (TN), respectively. True diploids in the putative haploid fraction and true haploids in the putative diploid fraction that were also confirmed by the vigor test and cytometry were categorized as false positives (FP) and false negatives (FN), respectively. The numbers of TP, TN, FP, and FN were adjusted in each population based on the total number of putative haploids and diploids identified in the population based on the *R1-nj*. The data resulting from the level of ploidy classification were used to calculate the false positive rate (FPR), false negative rate (FNR), and Matthews correlation coefficient (MCC), according to the following equations: 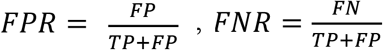, and 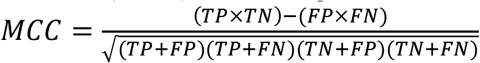, where FPR is the proportion of diploids in the seed fraction classified as putative haploids, FNR is the proportion of true haploids misclassified as diploid seeds, and MCC indicates the correlation between the actual and predicted ploidy level based on the marker *R1-nj*. The MCC has been suggested as a balanced measure of the quality of binary classifications, especially when the binary classes are of substantially different sizes (Chaikam et al., 2016). The MCC was developed by Brian W. Matthews and has become essential for measuring the performance of binary classification models. This coefficient fluctuates within the interval (−1, 1), and the higher its value is, the better the prediction ability of the classifier. On the other hand, when the assumed value is 0, there is no correlation between the two variables; i.e., the model is predicting randomly (Matthews, 1975).

For all characteristics, we assumed *Y*_*ij*_ ∼ *binomial*(*m*_*ij*_, *π*_*ij*_), where *Y*_*ij*_ is the total number of seeds of the *i*-th genotype in replication *j, m*_*ij*_ is the number of putative haploid seeds, and *π*_*ij*_ is the probability of success (probability of putative haploid seeds). We fit a generalized linear model (GLM) with a logit link function described by the following linear predictor:

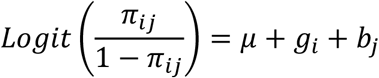

To evaluate the goodness of fit of the model, we used visual inspection using seminormal plots, using the hnp package (Moral et al., 2017). To compare the evaluated genotypes, we used asymptotic confidence intervals obtained by the emmeans package (Russell, 2020). All statistical analyses were performed in the R environment (Team, 2021).

## Results

### Seed produced by crossing KHI x popcorn

Of the ten inbreed lines pollinated by KHI, five did not produce seeds and were not considered in the analysis. The inbreed lines 19-1206-6, 19-1504-4, 18-390-1, and 18-303-2, for sure because they are derived from temperate germplasm, present unilateral cross-incompatibility alleles. Although tropical, the inbred line 20-2039 also did not produce seeds, probably because it contains these same alleles. Of the crosses that were fertile, a total of 63,232 seeds were obtained from 10 temperate (9,506), six tropical (44,019), and four tropical x temperate (19,707) germplasm sources. The average number of seeds/ear in temperate germplasm was 76, lower than that found in tropical germplasm (98).

### Haploid/diploid misclassification using the Navajo phenotype

The FPR using the evaluation of seedling vigor as the “gold standard” ranged from 0 to 65.4%, with a mean of 26.2% (Table 2). Among the popcorn females, 65% presented FPR > 20%. The MCC value ranged from 0.32 to 1, with a mean of 0.77. The average FPR found in the inbred lines (28.5%) and interpopulation hybrids (26.3%) were slightly higher than the average value found for the populations (25.0%). The average FPR value found in the temperate germplasm (18.6%) was approximately 47% lower than that found in the tropical germplasm (35.0%). In the tropical x temperate germplasm, the average FPR value was 20.4%. Among the putative haploids generated from 20-2009, UFV-MP1, UFV-MP4, UFV-MP2 x UFV-MP5, and Synthetic UFV x UFV-MP1, no false positive was identified and the MCC was 1. Among the inbred lines, the genotype 18-176-5 had the highest FPR (65.7%) but high estimates were also found for females Beija-Flor C4 and Beija-Flor C4 x Synthetic UFV (FPR > 50%). The average FPR found in the controls and popcorn were similar, 26.3 and 25.6%, respectively. There were no false negatives. However, the sample of putative diploids evaluated may have been.

**Table 2.**
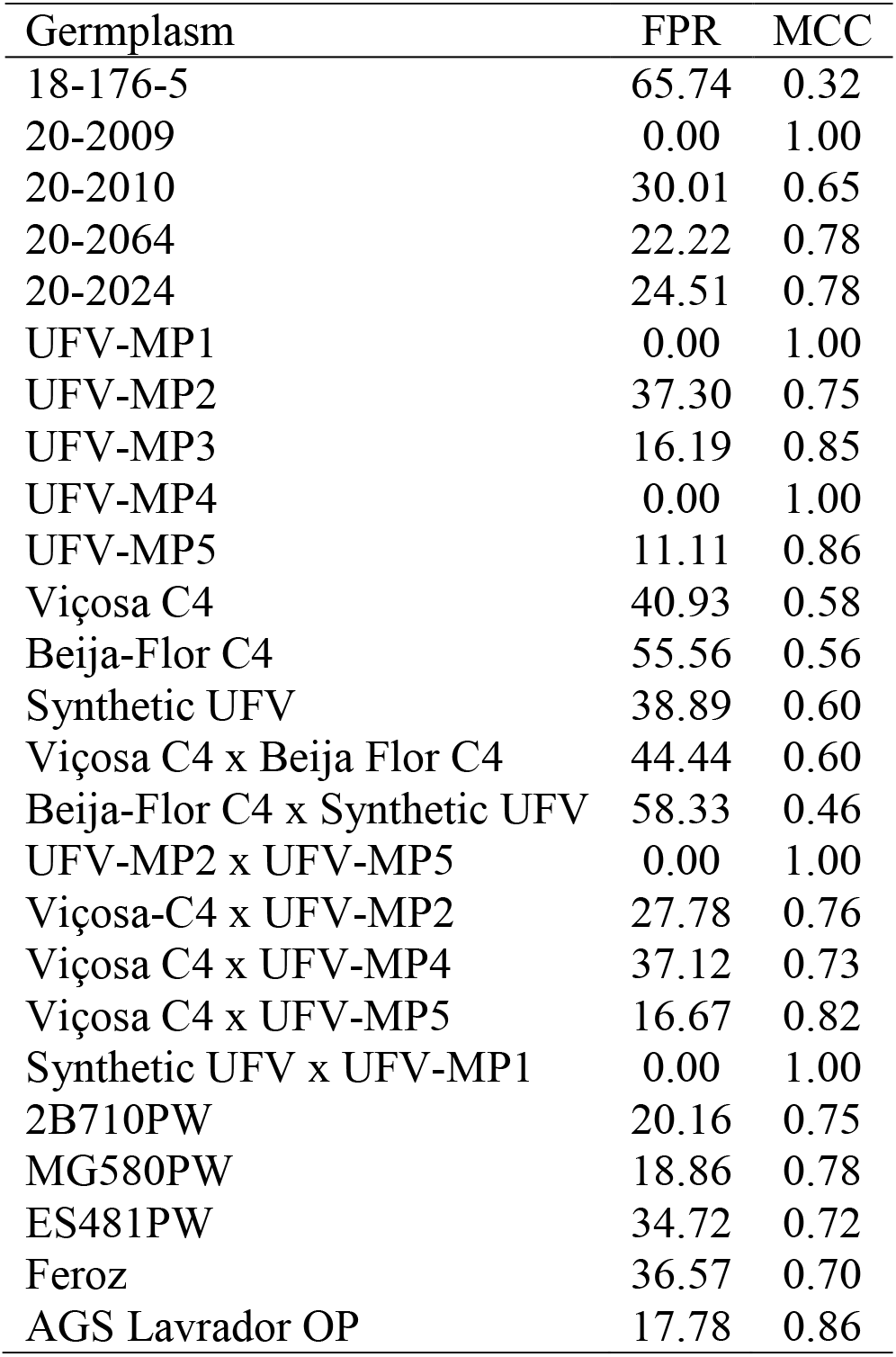
False positive rate and Matthews correlation coefficient based on “gold standard” vigor.

The FPR considering the level of ploidy determined by the flow cytometry as the “gold standard”, ranged from 0 to 100% among the 14 germplasm sources evaluated. The average FPR obtained by the Navajo phenotype and by the Navajo phenotype associated with subsequent elimination of identified false positive by vigor was 76.4 and 69.2%, respectively (Table 3). The inbred lines had the lowest mean FPR (60.6%) compared to the populations (88.5%) and interpopulation hybrids (82.6%). Most of the studied females showed high FPR values (FPR > 50%). The elimination of false positives by the vigor test reduced the FPR for most of the evaluated females, except for females 18-176-5, 20-2009, UFV-MP1, Beija-Flor C4 and Synthetic UFV x UFV-MP1. The average MCC values were low (0.38). Again, no false negatives were identified.

**Table 3.** False positive rate and Matthews correlation coefficient of Navajo phenotype and Navajo phenotype after elimination of false positives by vigor, using the ploidy level as the “gold standard”.

| Germplasm | Navajo phenotype |  | Navajo phenotype and vigor |  |
| --- | --- | --- | --- | --- |
|  | FPR | MCC | FPR | MCC |
| 18-176-5 | 100.00 | 0.00 | 100.00 | 0.00 |
| 20-2009 | 8.33 | 0.93 | 8.33 | 0.93 |
| 20-2010 | 85.78 | 0.18 | 89.96 | 0.25 |
| 20-2064 | 33.92 | 0.58 | 50.00 | 0.72 |
| 20-2024 | 38.59 | 0.57 | 54.58 | 0.71 |
| UFV-MP1 | 66.67 | 0.68 | 66.67 | 0.68 |
| UFV-MP5 | 83.33 | 0.42 | 88.89 | 0.48 |
| Viçosa C4 | 90.49 | 0.14 | 94.43 | 0.22 |
| Beija-Flor C4 | 100.00 | 0.00 | 100.00 | 0.00 |
| Synthetic UFV | 87.00 | 0.17 | 92.59 | 0.25 |
| Viçosa-C4 x UFV-MP2 | 58.33 | 0.42 | 77.78 | 0.54 |
| Viçosa C4 x UFV-MP4 | 46.83 | 0.39 | 70.08 | 0.52 |
| Synthetic UFV x UFV-MP1 | 100.00 | 0.00 | 100.00 | 0.00 |
| MG580PW | 59.35 | 0.39 | 67.11 | 0.47 |

### Induction rate

Based on the binomial analysis of the HIR, which was adjusted after the elimination of false positives identified through the vigor test, a significant effect was found for germplasm sources (p < 0.01). The mean HIR for the KHI inducer in the evaluated germplasm sources was 0.76% (Table 4). Of the 677 putative haploid seedlings, 459 were confirmed as true haploid, being 126 of maize, and 333 of popcorn (Supplementary Table 3).The overall average HIR of the KHI inducer in popcorn was 0.93%; while in the maize the overall average value was much lower (0.07%). The HIR in the tropical popcorn germplasm was 1.5 times lower than in temperate germplasm, with means of 0.89 and 1.32%, respectively. The average induction among the inbred lines was 1.57%, while the average in the populations was 0.85%. The lowest average induction was found in the interpopulation hybrids (0.57%). Of the populations, UFV-MP4 had the highest mean induction. However, the confidence interval was large, with lower and upper limits between 0.76 and 7.07%.

**Table 4.** Haploid induction rate adjusted by vigor and the confidence interval (CI) at 95% probability.

| Germplasm | HIR | CI |
| --- | --- | --- |
| 18-176-5 | 1.41 | 1.03-1.95 |
| 20-2009 | 1.41 | 0.74-2.69 |
| 20-2010 | 1.32 | 0.98-1.77 |
| 20-2064 | 1.86 | 1.24-2.78 |
| 20-2024 | 1.86 | 1.23-2.81 |
| UFV-MP1 | 0.44 | 0.17-1.18 |
| UFV-MP2 | 0.31 | 0.15-0.62 |
| UFV-MP3 | 1.76 | 1.00-3.07 |
| UFV-MP4 | 2.36 | 0.76-7.07 |
| UFV-MP5 | 0.60 | 0.22-1.58 |
| Viçosa C4 | 0.72 | 0.51-1.01 |
| Beija-Flor C4 | 0.37 | 0.12-1.15 |
| Synthetic UFV | 0.19 | 0.13-0.27 |
| Viçosa C4 x Beija Flor C4 | 0.10 | 0.06-0.31 |
| Beija-Flor C4 x Synthetic UFV | 0.21 | 0.02-1.04 |
| UFV-MP2 x UFV-MP5 | 2.32 | 1.85-3.03 |
| Viçosa-C4 x UFV-MP2 | 0.25 | 0.11-0.55 |
| Viçosa C4 x UFV-MP4 | 0.20 | 0.12-0.33 |
| Viçosa C4 x UFV-MP5 | 0.06 | 0.02-0.19 |
| Synthetic UFV x UFV-MP1 | 0.85 | 0.72-1.05 |
| 2B710PW | 0.08 | 0.06-0.11 |
| MG580PW | 0.08 | 0.06-0.10 |
| ES481PW | 0.03 | 0.02-0.05 |
| Feroz | 0.11 | 0.06-0.18 |
| AGS Lavrador OP | 0.04 | 0.02-0.07 |

Based on the binomial analysis of the HIR, adjusted after the elimination of the false positives identified by the ploidy level, a significant effect was found for the germplasm sources (p < 0.01). The overall mean HIR for the KHI inducer in the 14 germplasm sources evaluated was 0.32% (Table 5). Of the 677 putative haploid seedlings evaluated, only 78 were confirmed as true haploid by ploidy level (Supplementary Figure 1), being 18 of maize, and 60 of popcorn (Supplementary Table 4). After adjusting the means by the FPR estimated through the FP identified by the ploidy level, the mean HIR of the KHI inducer in popcorn was 0.34%. The HIR found for the inbred lines, populations, and interpopulation hybrids were 0.75, 0.09, and 0.07%, respectively. Only the genotypes 20-2009, 20-2064 and 20-2024 presented HIR > 1%; in the other evaluated germplasm sources, the HIR values were lower than 0.25%.

**Table 5.**
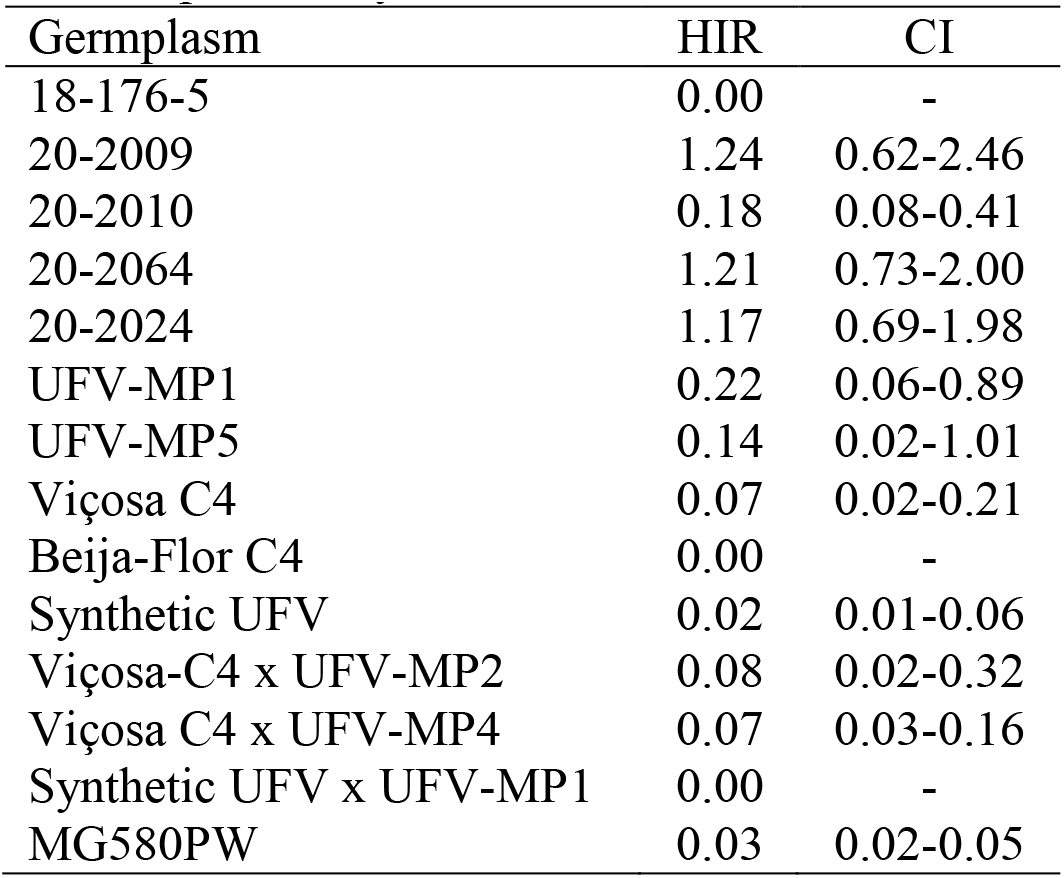
Haploid induction rate adjusted by the ploidy level and the confidence interval (CI) at 95% probability.

### Penetrance of the *R1-nj*

Binomial analysis of *R1-nj* penetrance showed significant effects for germplasm sources (p < 0.01). The mean penetrance between the evaluated germplasm sources was 71.9% (Table 6). The mean penetrance of popcorn (69.9%) was lower than the mean of the control (79.9%). Seventy percent of the evaluated popcorn materials had penetrance < 65%, 40% had values between 65 and 80%, and 30% had penetrance > 80%. The mean value of *R1-nj* penetrance in germplasms of tropical origin (67.2%) was lower than that of germplasms of origin (tropical x temperate) (67.8%) and temperate (74.6%). The inbred lines had the highest mean penetrance (80.2%), followed by populations (69.0%) and interpopulation hybrids (63.2%). The population Beija-Flor C4 presented the lowest penetrance value (30.8%), while genotype 20-2009 presented the highest value (98.6%).

**Table 6.**
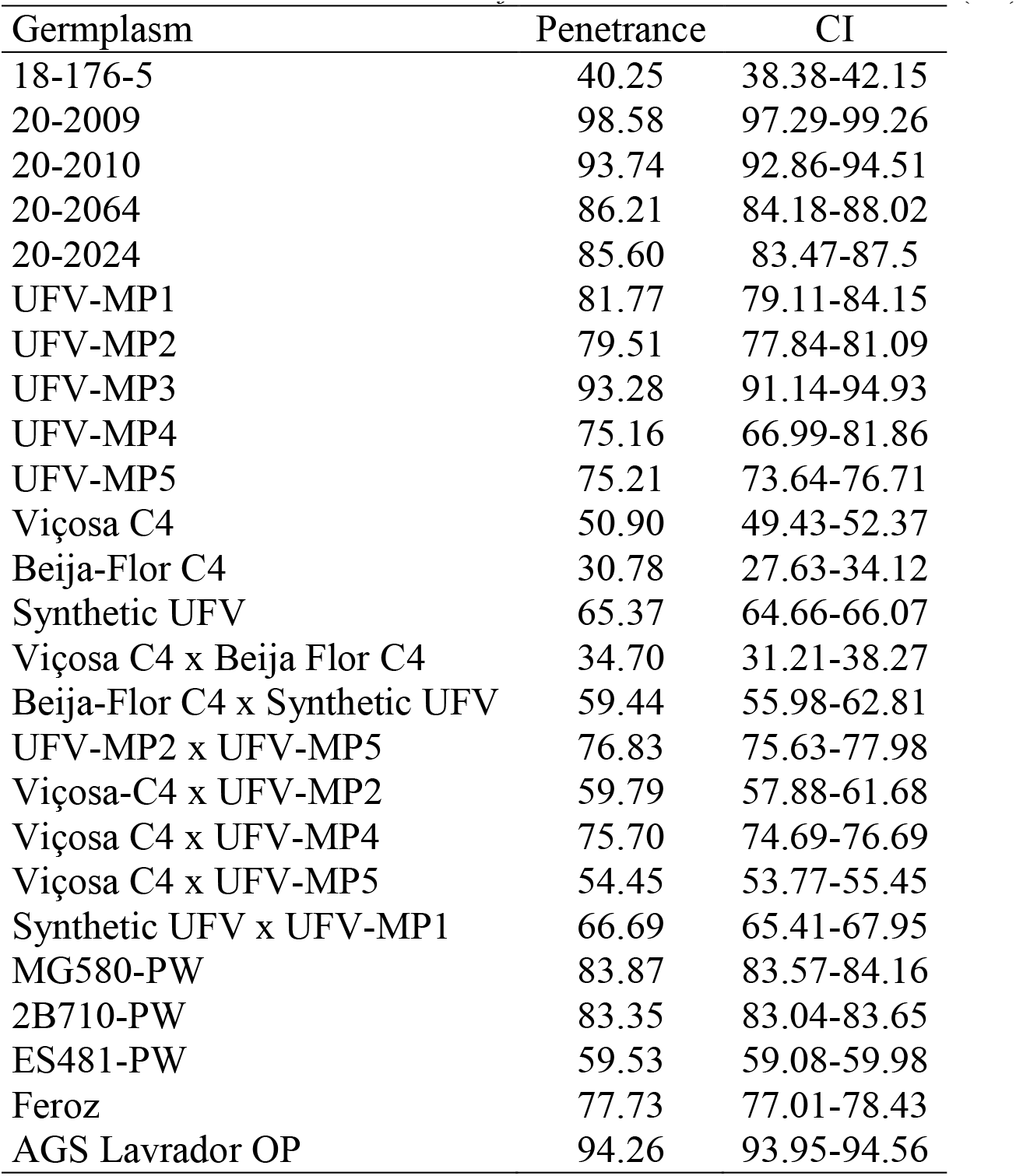
Penetrance of the *R1-nj* and the confidence interval (CI) at 95% probability.

| Germplasm | Penetrance | CI |
| --- | --- | --- |
| 18-176-5 | 40.25 | 38.38-42.15 |
| 20-2009 | 98.58 | 97.29-99.26 |
| 20-2010 | 93.74 | 92.86-94.51 |
| 20-2064 | 86.21 | 84.18-88.02 |
| 20-2024 | 85.60 | 83.47-87.5 |
| UFV-MP1 | 81.77 | 79.11-84.15 |
| UFV-MP2 | 79.51 | 77.84-81.09 |
| UFV-MP3 | 93.28 | 91.14-94.93 |
| UFV-MP4 | 75.16 | 66.99-81.86 |
| UFV-MP5 | 75.21 | 73.64-76.71 |
| Viçosa C4 | 50.90 | 49.43-52.37 |
| Beija-Flor C4 | 30.78 | 27.63-34.12 |
| Synthetic UFV | 65.37 | 64.66-66.07 |
| Viçosa C4 x Beija Flor C4 | 34.70 | 31.21-38.27 |
| Beija-Flor C4 x Synthetic UFV | 59.44 | 55.98-62.81 |
| UFV-MP2 x UFV-MP5 | 76.83 | 75.63-77.98 |
| Viçosa-C4 x UFV-MP2 | 59.79 | 57.88-61.68 |
| Viçosa C4 x UFV-MP4 | 75.70 | 74.69-76.69 |
| Viçosa C4 x UFV-MP5 | 54.45 | 53.77-55.45 |
| Synthetic UFV x UFV-MP1 | 66.69 | 65.41-67.95 |
| MG580-PW | 83.87 | 83.57-84.16 |
| 2B710-PW | 83.35 | 83.04-83.65 |
| ES481-PW | 59.53 | 59.08-59.98 |
| Feroz | 77.73 | 77.01-78.43 |
| AGS Lavrador OP | 94.26 | 93.95-94.56 |

## Discussion

For the first time, we induced haploids *in vivo* in popcorn. We observed induction in tropical, temperate, and tropical x temperate germplasm. However, the number of seeds obtained per ear from crossing the KHI inducer with germplasm sources of temperate origin was much lower than that obtained from tropical germplasm. The temperate germplasm used for the induction of haploids is of North American origin, which would lead us to assume that they would be fixed for the incompatibility allele *Ga1-s* (Kermicle and Evans, 2010). However, as there was seed production, it is more likely that they are *Ga1-s/ga1-s*, as this allele is codominant in relation to the *ga1* allele, that is, the *Ga1-s/ga1-s* genotypes have partial fertility for *ga1* pollen. Another hypothesis to explain the partial fertility observed in temperate germplasm is that the inducer is segregating. However, four inbred lines of temperate origin did not produce seeds, reducing the evidence for this hypothesis. Furthermore, these four genotypes are likely to be *Ga1-s/Ga1-s*, 100% incompatible with *ga1* pollen. As exposed, there is evidence that the KHI inducer is *ga1/ga1*. In addition, it is likely that most inducers used for induction of haploids in maize have this genotypic constitution, which would prevent *in vivo* induction by gymnogenetic inducers in sources of *Ga1-s/Ga1-s* germplasm, the most common genotype in popcorn (Kermicle and Evans, 2010). In contrast, temperate germplasm is the most targeted by popcorn breeders. We advise two viable ways to overcome the inducer/popcorn incompatibility barrier: employing androgenetic inducers or modifying the genotypic constitution of gymnogenetic inducers. However, androgenetic inducers are not as improved for HIR as gymnogenetic inducers. Additionally, it is not possible to use isolated fields, that is, pollination has to be done manually. Therefore, we recommend the introgression of alleles that overcome the fertility barrier imposed by *Ga1-s*, such as the *Ga1-s* allele itself or the *Ga1-m* allele.

Concerning the efficacy of the Navajo phenotype, we expected to find higher FPR in popcorn due to the smaller size and rounded shape of the seeds that make it difficult to visualize the embryo. However, the average FPR found in popcorn was lower than that found in the checks. The FPR values found in tropical and tropical x temperate germplasms were higher than that found in temperate germplasm. The high FPR associated with classification by the Navajo phenotype can be attributed to the presence of inhibitory genes, varied expressivity, and incomplete penetrance of this marker (Belicuas et al., 2007). In addition to environmental factors, such as the moisture content of the seeds and air pockets under the pericarp, these genes can also affect the color intensity and consequently lead to an erroneous classification of the ploidy level of the evaluated classes (Rotarenco et al., 2009; Prigge et al., 2011). Although classification based on Navajo phenotype leads to high magnitude FPR, FNR values were zero. This is especially important for inducers with low HIR, as is the case with the KHI inducer, because even with the increase in the size of plant populations in the induction fields, as recommended by Melchinger et al., (2013), there are a limited number of available haploids.

Elimination of false positives by vigor assessment reduced the FPR for most females evaluated. However, the mean values of MCC remained of low magnitude, indicating a low correlation between the ploidy determined by *R1-nj* and the true level of ploidy of the evaluated seedlings. Melchinger et al., (2013) found high FPR values, ranging from 7.8 to 67.6%, using the *R1-nj* marker. The elimination of false positives through the evaluation of seedling vigor can be useful, as it is an easy and low-tech method that allows reliably eliminating a fraction of the false positives, since none of the 218 false positives identified was classified as true haploid by the ploidy level. Eliminating false positives before chromosome duplication is an important strategy to reduce cost with reagents, waste of space, inputs in greenhouses and/or DH_0_ seed multiplication field, and labor resources (Choe et al., 2012; Chaikam et al., 2015).

The KHI haploidy inducer showed an average HIR of 0.93% in popcorn after adjusting the FPR by the vigor and 0.34% if adjusted by the ploidy level. These rates are very low, considering that inducers with HIR > 10% are currently available, as in RWS and UH400 (Prigge et al., 2012; Chaikam et al., 2019). Although the HIR depends on both the inducer and the source of germplasm, the inducer has a greater contribution in determining this characteristic (Kebede et al., 2011). Thus, it is not possible to attribute the low HIR values found to popcorn.

To evaluate the expression of the *R1-nj* gene, we defined a variable obtained by the proportion between seeds that present the Navajo phenotype and the total number of induced seeds, and we called it penetrance. We found incomplete penetrance of the *R1-nj* in the popcorn germplasm as well as in the controls. This was expected, as one of the main limitations of *R1-nj* in haploid/diploid selection is its incomplete penetrance and varied expressivity. The low values of the penetrance estimates in tropical and tropical x temperate germplasms indicate a higher frequency of inhibitors in these germplasms, such as *C1-I, C2-Idf*, and *in-1D* (Chaikam et al., 2015; Gain et al., 2022). Chaikam et al., (2015) propose a careful selection of the parents that will be used in the synthesis of the source germplasm, regarding the absence of anthocyanin inhibitors. The lower penetrance estimate of *R1-nj* in tropical popcorn germplasm indicates the need to use inducers equipped with other haploid/diploid classification systems that can complement *R1-nj* to identify haploids more efficiently. The average penetrance showed by the inbred lines was high. This result is important since the most used strategy for the production of DH lines is the induction of haploids from bi-parental crosses between elite lines (Chaikam et al., 2019).

## Conclusions

There is induction of haploids in germplasm of tropical and temperate origin. We recommend the selection of haploids associating the Navajo phenotype with a direct method of confirming the ploidy level, such as flow cytometry. We also show that haploid screening based on Navajo phenotype and seedling vigor reduces misclassification. The origin and genetic background of the source germplasm influence the *R1-nj* penetrance. Because the known inducers are maize, developing doubled-haploid technology for popcorn hybrid breeding requires overcoming the unilateral cross-incompatibility.

## Abbreviations

DH: Doubled-haploid;
FPR: False positive rate;
FN: False negative;
FNR: False negative rate;
FP: False positive;
HIR: Haploid induction rate;
R*1-nj*: R*1-navajo* gene marker;
MCC: Matthews correlation coefficient;
TN: True negative;
TP: True positive.

## Author Contributions

JM and JP contributed to the project planning and overall coordination. JP performed and coordinated the field experiments. JC, WR, JP and VT were responsible for the flow cytometry. KO, JM and JP carried out the analysis. JP wrote the manuscript. All authors have made their contribution to editing the manuscript and approved the final version.

## Acknowledgments

I thank the National Council for Scientific and Technological Development (CNPq), the Brazilian Federal Agency for Support and Evaluation of Graduate Education (Capes; Finance Code 001), and the Foundation for Research Support of Minas Gerais State (Fapemig) for financial support.

